# Diversity of Antimicrobial-Resistant *Aeromonas* Species Isolated from Aquatic Environments in Brazil

**DOI:** 10.1101/2020.06.23.168385

**Authors:** Danieli Conte, Jussara Kasuko Palmeiro, Adriane de Almeida Bavaroski, Luiza Souza Rodrigues, Daiane Cardozo, Ana Paula Tomaz, Josué Oliveira Camargo, Libera Maria Dalla-Costa

## Abstract

In the present study, we characterized antimicrobial resistance profile and genetic relatedness of *Aeromonas* spp. isolated from healthcare and urban effluents, wastewater treatment plant (WWTP), and river water. We detected the presence of genes responsible for the resistance to β-lactam, quinolone, and aminoglycoside. Enterobacterial Repetitive Intergenic Consensus PCR and multilocus sequence typing (MLST) were carried out to differentiate the strains and multilocus phylogenetic analysis (MLPA) was used to identify species. A total of 28 *Aeromonas* spp. cefotaxime-resistant strains were identified that carried a variety of resistance determinants, including uncommon GES-type β-lactamases. Multidrug-resistant *Aeromonas* spp. were found in hospital wastewater, WWTP, and sanitary effluent. Among these isolates, we detected *A. caviae* producing GES-1 or GES-5, as well as *A. veronii* harboring GES-7 or GES-16. We successfully identified *Aeromonas* spp. by using MLPA and found that *A. caviae* was the most prevalent species (85.7%). In contrast, it was not possible to determine sequence type of all isolates, suggesting incompleteness of the *Aeromonas* spp. MLST database. Our findings reinforce the notion about the ability of *Aeromonas* spp. to acquire determinants of antimicrobial resistance from the environment. Such ability can be enhanced by the release of untreated healthcare effluents, in addition to the presence of antimicrobials, recognized as potential factors for the spread of resistance. Thus, *Aeromonas* spp. could be included as priority pathogens under the One Health concept.

**IMPORTANCE:** *Aeromonas* species are native bacteria in aquatic ecosystems worldwide. However, they have also been isolated from humans and animals. Globally, aquatic environments have been affected by anthropogenic activities. For example, the excessive use of antimicrobials in medical and veterinary practice causes the development of bacterial resistance. In addition, eliminated hospital and sanitary effluents can also serve as potential sources of bacteria carrying antimicrobial resistance genes. Thereby, impacted environments play an important role in the transmission of these pathogens, their evolution, and dissemination of genes conferring resistance to antimicrobials. *Aeromonas* spp. have been reported as a reservoir of antimicrobial resistance genes in the environment. In this study, we identified a great repertoire of antimicrobial resistance genes in *Aeromonas* spp. from diverse aquatic ecosystems, including those that encode enzymes degrading broad-spectrum antimicrobials widely used to treat healthcare-associated infections. These are a public health threat as they may spread in the population.

## INTRODUCTION

Members of the genus *Aeromonas* are food and waterborne opportunistic pathogens in humans and animals (1). As human pathogens, they can cause gastrointestinal disease and serious life-threatening extraintestinal infections, such as wound and skin soft-tissue infections, liver abscesses, bacteremia, and meningitis in both immunocompromised and healthy hosts (2, 3).

It is already consensus that the widespread and often inappropriate use of antimicrobials accelerates the emergence of resistant bacterial pathogens (4). The selective pressure exerted by these chemical compounds becomes even more concerning when considering their impact on the environment, and not only in the restricted settings, such as hospitals. In the environment, these compounds can accumulate and circulate between different communities, increasing the environmental resistome and creating reservoirs of new resistance genes (5). In this context, *Aeromonas* spp., widely distributed in the soil and aquatic ecosystems, have played an important role as a reservoir of antimicrobial resistance genes (ARG) (6–8).

*Aeromonas* spp. can harbor at least three chromosomal β-lactamases of Ambler classes B, C, and D, which may confer resistance to penicillins, cephalosporins, and carbapenems (9). Although some species can exhibit intrinsic resistance to β-lactam by unrepressed gene expression, most of them remain susceptible to aminoglycosides, chloramphenicol, tetracycline, trimethoprim-sulfamethoxazole, and quinolones (2).

Through the natural transformation, *Aeromonas* spp. are able to acquire mobile genetic elements harboring antimicrobial resistance determinants (10–12). For example, plasmid-mediated quinolone resistance (e.g., *qnr*S and *aac(6′)-Ib-cr*), carbapenemase (e.g., *bla*_KPC,_ *bla*_NDM_, *bla*_GES_, *bla*_IMP_, *bla*_VIM,_ and *bla*_OXA-48_), and aminoglycosides-encoding genes (e.g., *rmt*D) have been detected in *Aeromonas* spp. from water sources (13–22).

Several studies aimed to understand the phylogenetic relationship of the genus *Aeromonas*; however, its taxonomy is still a challenge due to complex classification of this genus and continuous additions of new species (9, 23). Genotyping techniques using sequences of housekeeping genes, such as multilocus sequence typing (MLST) and multilocus phylogenetic analysis (MLPA), have helped to identify many of the new species of this genus by overcoming the limitation of the high similarity of the 16S rRNA gene among the closely related species. These methods have been applied to different bacterial genera for species identification and ecological studies (24, 25).

The interest in this genus has grown over the past two decades due to (i) worldwide distribution of aeromonads, (ii) challenges to accurate identification and classification of different species, (iii) occurrence of strains with antimicrobial resistance, including resistance to carbapenem, and (iv) ability of some strains to survive conventional wastewater treatments (18, 26, 27). In this study, we evaluated determinants of resistance to β-lactam, quinolone, and aminoglycoside in *Aeromonas* spp. isolated from different aquatic environments. Further, we performed a detailed phylogenetic analysis to identify species and lineages associated with the spread of antimicrobial resistance.

## RESULTS

### Screening for Aeromonads in aquatic environments and determination of their antimicrobial susceptibility profiles

A total of 158 colonies was screened. Ninety colonies (57%, n = 90/158) were presumptively screened as *Aeromonas* spp. based on oxidase and glucose fermentation tests. Among these, 45 isolates were identified as *Aeromonas* spp. by VITEK^®^ 2 System and mass spectrometry [matrix assisted laser desorption ionization-time of flight mass spectrometry (MALDI-TOF)]. After Enterobacterial Repetitive Intergenic Consensus (ERIC)-PCR analysis, 42 unique isolates were evaluated for antimicrobial susceptibility profile. Out of them, 28 cefotaxime-resistant isolates were further studied.

Table 1 summarizes the results of antimicrobial susceptibility testing, phenotypic tests for extended-spectrum β-lactmases (ESBLs) and carbapenemases, and detection of resistance genes according to sample isolation sites. Nearly 90% of isolates had ESBL phenotype (n = 25/28), however some isolates showed susceptibility to ceftazidime (28.6%, n = 8) and cefepime (57.1 %, n = 16). ESBL phenotypic test poorly detected GES-type (isolates 4 and 11). Almost all environmental samples were found to be resistant to gentamicin (61%, n = 17) and ciprofloxacin (32%, n = 9), except for those from river water [Figure 1, upstream river water (URW) and downstream river water (DRW)]. Few isolates displayed resistance to three or more distinct classes of antimicrobials and were classified as multidrug-resistant (MDR) (28). These isolates were detected in the hospital and sanitary effluents, as well as in WWTP (isolates 2, 6, 11, 16, 28).

**TABLE 1.** Antimicrobial resistance profile of *Aeromonas* spp. isolated from diverse aquatic environments HPP, Pequeno Principe Hospital; ECU, Emergency Care Unit; CHC-UFPR, Complex Clinics Hospital of the Federal University of Paraná; WWTP, wastewater treatment plant; RW, river water; SE, sanitary effluent; ND, not determined; CAZ, ceftazidime; FEP, cefepime; CTX, cefotaxime; CIP, ciprofloxacin; GNT, gentamicin; ERT, ertapenem; MER, meropenem; IMI, imipenem.

| Sample collection site | Isolates ID | Species | Minimal inhibitory concentration (µg/mL) |  |  |  |  |  |  |  | ESBL | Carbapenemases |  | Resistance genes |
| --- | --- | --- | --- | --- | --- | --- | --- | --- | --- | --- | --- | --- | --- | --- |
|  |  |  | CAZ | FEP | CTX | CIP | GNT | ERT | MER | IMI |  | class A | class B |  |
| HPP | 1 | *A. caviae* | 2 | 16 | > 32 | 2 | > 8 | 0.0312 | 0.0312 | 0.125 | + | — | — | *bla*<sub>CTX-M-1</sub>, *bla*<sub>TEM</sub> |
|  | 2 | *A. caviae* | > 16 | 32 | > 32 | 4 | > 8 | 0.0312 | 0.0312 | 0.125 | + | — | — | *bla*<sub>TEM</sub>, *aac*(6')-Ib-cr |
|  | 3 | *A. caviae* | 1 | < 4 | 32 | 2 | > 8 | 0.0312 | 0.0312 | 0.125 | + | — | — | *bla*<sub>CTX-M-1</sub>, *bla*<sub>TEM</sub> |
|  | 4 | *A. caviae* | 8 | < 4 | 4 | < 0.5 | < 2 | 0.0312 | 0.0312 | 0.125 | — | — | — | *bla*<sub>GES-1</sub> |
|  | 5 | *A. caviae* | 8 | < 4 | 4 | < 0.5 | < 2 | 0.0312 | 0.0312 | 0.125 | + | — | — | *bla*<sub>GES-1</sub> |
|  | 6 | *A. veronii* | > 16 | 32 | > 32 | < 0.5 | > 8 | > 8 | > 8 | > 128 | + | + | — | *bla*<sub>GES-16</sub> |
|  | 7 | *A. caviae* | > 16 | 32 | > 32 | 2 | > 8 | 0.0156 | 0.0312 | 0.125 | + | — | — | *bla*<sub>TEM</sub> |
|  | 8 | *A. caviae* | > 16 | 32 | > 32 | 2 | > 8 | 0.0156 | 0.0312 | 0.125 | + | — | — | *bla*<sub>TEM</sub> |
|  | 9 | *A. caviae* | 4 | 8 | 4 | 4 | 4 | 1 | 0.125 | 0.125 | — | — | — | *aac*(6')-Ib-cr |
|  | 10 | *A. caviae* | > 16 | 16 | > 32 | 2 | > 8 | 0.0156 | 0.0312 | 0.125 | + | — | — | *bla*<sub>TEM</sub> |
|  | 11 | *A. veronii* | > 16 | 16 | 64 | 4 | > 8 | > 8 | > 8 | > 128 | — | — | + | *bla*<sub>GES-7</sub>, *aac*(6')-Ib-cr |
| ECU | 12 | *A. caviae* | 2 | 8 | 64 | < 0.5 | 8 | 0.0156 | 0.0312 | 0.125 | + | — | — | *bla*<sub>CTX-M-1</sub> |
| CHC-UFPR | 13 | *A. caviae* | 4 | 32 | > 32 | < 0.5 | > 8 | < 0.25 | < 0.5 | < 0.5 | + | — | — | *bla*<sub>CTX-M-1</sub>, *bla*<sub>CTX-M-2</sub> |
|  | 14 | *A. caviae* | < 1 | < 4 | > 32 | > 2 | > 8 | < 0.25 | < 0.5 | < 0.5 | + | — | — | *bla*<sub>CTX-M-1</sub>, *bla*<sub>TEM</sub> |
|  | 15 | *A. caviae* | < 1 | < 4 | > 32 | > 2 | > 8 | < 0.25 | < 0.5 | < 0.5 | + | — | — | *bla*<sub>CTX-M-1</sub>, *bla*<sub>TEM</sub>, *aac*(6')-Ib-cr, *qnrS* |
|  | 16 | *A. caviae* | 8 | 64 | > 32 | > 2 | > 8 | > 8 | > 8 | 8 | + | + | — | *bla*<sub>GES-5</sub> |
|  | 17 | *A. caviae* | > 16 | < 4 | > 32 | 2 | > 8 | < 0.25 | < 0.5 | < 0.5 | + | — | — | ND |
|  | 18 | *A. caviae* | 16 | < 4 | > 32 | 2 | > 8 | < 0.25 | < 0.5 | < 0.5 | + | — | — | ND |
| WWTP | 19 | *A. caviae* | 16 | > 32 | > 32 | > 2 | > 8 | < 0.25 | < 0.5 | < 0.5 | + | — | — | *bla*<sub>CTX-M-1</sub> |
|  | 20 | *A. caviae* | > 16 | < 4 | 32 | < 0.5 | < 2 | < 0.25 | < 0.5 | < 0.5 | + | — | — | *bla*<sub>CTX-M-8</sub>, *bla*<sub>CTX-M-9</sub> |
|  | 21 | *A. caviae* | 16 | < 4 | 32 | < 0.5 | < 2 | < 0.25 | < 0.5 | < 0.5 | + | — | — | *bla*<sub>CTX-M-8</sub>, *bla*<sub>CTX-M-9</sub> |
|  | 22 | *A. sanarellii* | > 16 | < 4 | > 32 | < 0.5 | < 2 | < 0.25 | < 0.5 | < 0.5 | + | — | — | ND |
|  | 23 | *A. caviae* | 16 | < 4 | 32 | < 0.5 | < 2 | < 0.25 | < 0.5 | < 0.5 | + | — | — | ND |
|  | 24 | *A. caviae* | > 16 | < 4 | > 32 | < 0.5 | < 2 | < 0.25 | < 0.5 | < 0.5 | + | — | — | ND |
| RW | 25 | *A. caviae* | > 16 | < 4 | 32 | < 0.5 | < 2 | < 0.25 | < 0.5 | < 0.5 | + | — | — | ND |
|  | 26 | *A. sanarellii* | < 1 | < 4 | > 32 | 2 | 8 | < 0.25 | < 0.5 | < 0.5 | + | — | — | *bla*<sub>CTX-M-1</sub>, *bla*<sub>TEM</sub> |
| SE | 27 | *A. caviae* | > 16 | 16 | > 32 | > 2 | > 8 | < 0.25 | < 0.5 | < 0.5 | + | — | — | *bla*<sub>TEM</sub>, *aac*(6')-Ib-cr, *qnrS* |
|  | 28 | *A. hydrophila* | 8 | 32 | > 32 | > 2 | > 8 | > 8 | > 8 | 4 | + | — | + | *bla*<sub>CTX-M-1</sub>, *aac*(6')-Ib-cr, *qnrS* |

**FIG 1.**
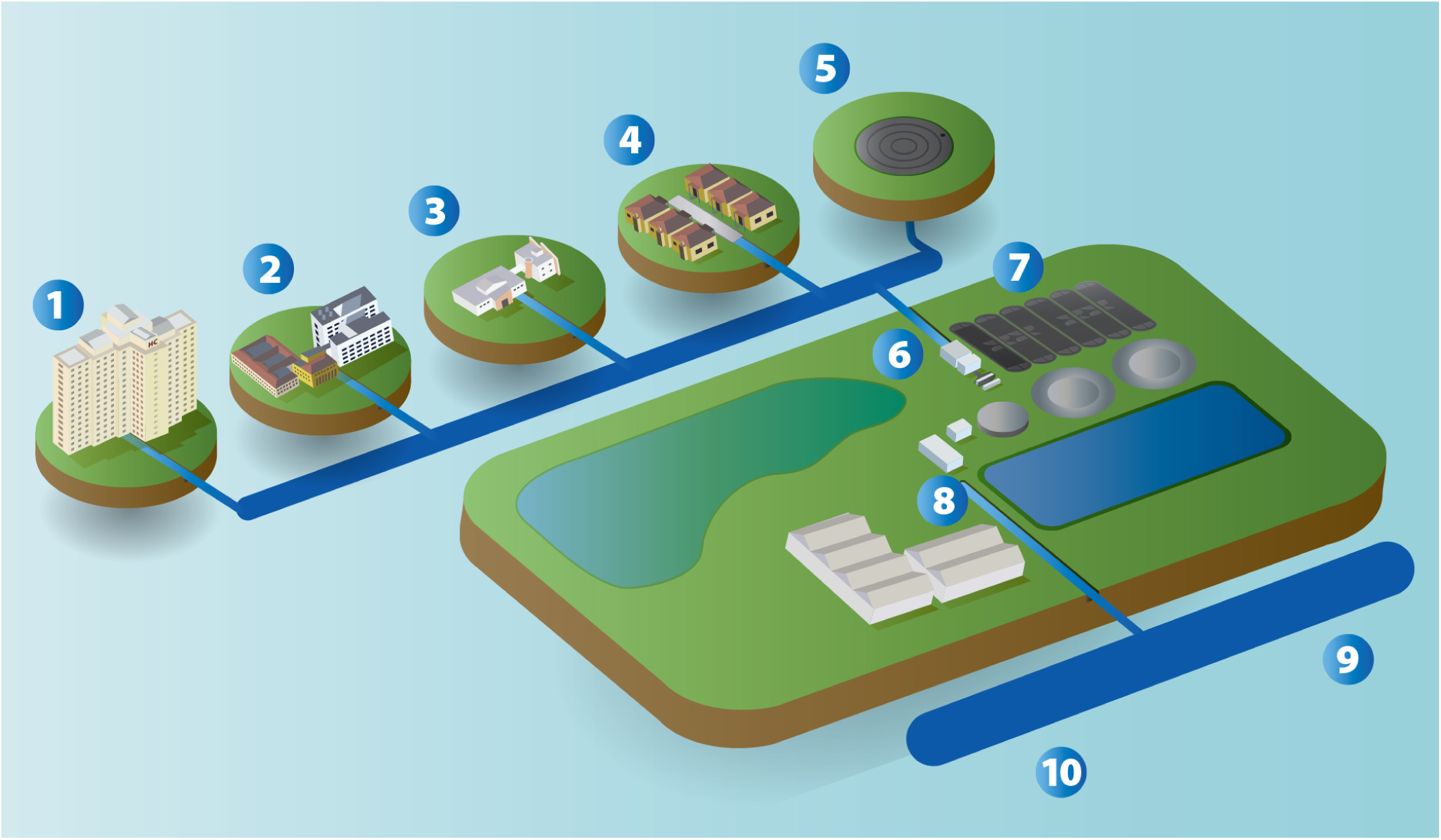
Graphical illustration of sample collection sites. (1) Complex Clinics Hospital of the Federal University of Paraná (CHC-UFPR); (2) Pequeno Principe Hospital (HPP); (3) emergency care unit (ECU); (4) domestic sewage; (5) sanitary effluent (SE); (6) inflow sewage; (7) aeration tank; (8) outflow sewage; (9) upstream river water (URW); (10) downstream river water (DRW). Parts 6, 7, and 8 correspond to the wastewater treatment plant (WWTP). CHC-UFPR is a 640-bed tertiary care teaching hospital. HPP is a 360-bed referral pediatric tertiary care hospital. WWTP serves 778,500 people. URW and DRW were sampled 100 m away from the point where treated WWTP was discharged.

ESBL-producing isolates included *bla*_CTX-M_ (39%, n = 11/28), *bla*_TEM_ (32%, n = 9/28), and *bla*_GES_ (10%, n = 3/28), with some co-production of different ESBL-type (Table 1). GES-type ESBL was identified in *A. caviae* (*bla*_GES-1_, isolates 4 and 5) and *A. veronii* (*bla*_GES-7_, isolate 11) from the hospital effluents. In addition, GES-type carbapenamase was detected in *A. caviae* (*bla*_GES-5_, isolate 16) and *A. veronii* (*bla*_GES-16_, isolate 6), also from the hospital effluents. Co-production of genes was rarely noted in GES-type producing isolates. *A. hydrophila* (isolate 28) and *A. veronii* (isolate 11) showed carbapenem resistance, but no carbapenemase genes were detected. Only five isolates had ESBL and plasmid-mediated quinolone resistance (PMQR) genes (isolates 2, 11, 15, 27, and 28). No 16S rRNA methyltransferases (16S RMTases) were identified.

### Phylogenetic analysis by MLPA, mass spectrometry, and automated phenotypic method

The phylogenetic tree allowed to identify all *Aeromonas* spp. isolates, revealing four different species, *A. caviae* (82.4%, n = 23), *A. veronii* (7%, n = 2), *A. sanarelli* (7%, n = 2), and *A. hydrophila* (3.6%, n = 1) (Figure 2). The convergence of the results between MLPA and MALDI-TOF Microflex LT Biotyper 3.0 occurred only for *A. veronii* and *A. hydrophila*. There was no agreement on the identification of *A. sanarelli*. In addition, 19 out of 23 *A. caviae* isolates were identified by MALDI-TOF Microflex LT Biotyper 3.0. VITEK^®^ 2 System and MALDI-TOF VITEK^®^ MS identified *Aeromonas* species with low accuracy (Table 2).

### Molecular typing by MLST

Among 28 *Aeromonas* isolates, only three showed match with all six alleles of housekeeping genes: *A. veronii* (isolate 11) assigned to ST 257 and *A. caviae* isolates 14 and 15 assigned to ST 94 (Figure 2). Three isolates had matches with five alleles, and MLST database indicated unique STs: ST 367 for *A. hydrophila* (isolate 28) and ST 584 for *A. caviae* (isolates 20 and 21). For the remaining isolates, MLST database was unable to define the STs.

**TABLE 2.** Comparison of results obtained by VITEK® 2, Microflex LT, and VITEK® MS instruments with phylogenetic analysis by MLPA for identifying *Aeromonas* spp. isolates

| Isolates ID | Automated colorimetric assay | Mass spectrometry |  | Phylogenetic analysis by MLPA |
| --- | --- | --- | --- | --- |
|  | VITEK® 2 System | Microflex LT Instrument | VITEK® MS Instrument |  |
| 1 | *A. hydrophila/caviae* | *A. caviae* | *A. hydrophila/puntacta* | *A. caviae* |
| 2 | *A. hydrophila/caviae* | *A. caviae/ hydrophila* | *A. hydrophila/puntacta* | *A. caviae* |
| 3 | *A. hydrophila/caviae* | *A. caviae* | *A. hydrophila/puntacta* | *A. caviae* |
| 4 | *A. hydrophila/caviae* | *A. caviae* | *A. hydrophila/puntacta* | *A. caviae* |
| 5 | *A. hydrophila/caviae* | *A. caviae/ hydrophila* | *A. hydrophila/puntacta* | *A. caviae* |
| 6 | *A. sobria* | *A. veronii* | *A. veronii/hydrophila/puntacta/sobria* | *A. veronii* |
| 7 | *A. hydrophila/caviae* | *A. caviae* | *A. hydrophila/puntacta* | *A. caviae* |
| 8 | *A. hydrophila/caviae* | *A. caviae* | *A. hydrophila/puntacta* | *A. caviae* |
| 9 | *A. hydrophila/caviae* | *A. caviae* | *A. hydrophila/puntacta* | *A. caviae* |
| 10 | *A. hydrophila/caviae* | *A. caviae* | *A. hydrophila/puntacta* | *A. caviae* |
| 11 | *A. hydrophila/caviae* | *A. veronii* | *A. hydrophila/puntacta/sobria* | *A. veronii* |
| 12 | *A. hydrophila/caviae* | *A. caviae* | *A. hydrophila/puntacta/sobria* | *A. caviae* |
| 13 | *A. hydrophila/caviae* | *A. caviae* | *A. hydrophila/puntacta* | *A. caviae* |
| 14 | *A. hydrophila/caviae* | *A. caviae* | *A. hydrophila/puntacta* | *A. caviae* |
| 15 | *A. hydrophila/caviae* | *A. caviae* | *A. hydrophila/puntacta* | *A. caviae* |
| 16 | *A. hydrophila/caviae* | *A. caviae* | *A. hydrophila/puntacta* | *A. caviae* |
| 17 | *V. cholerae* | *A. caviae* | *A. hydrophila/puntacta* | *A. caviae* |
| 18 | *A. hydrophila/caviae* | *A. caviae/ hydrophila* | *A. hydrophila/puntacta* | *A. caviae* |
| 19 | *Aeromonas spp.* | *A. caviae* | *A. hydrophila/puntacta* | *A. caviae* |
| 20 | *Aeromonas spp.* | *A. caviae* | *A. hydrophila/puntacta* | *A. caviae* |
| 21 | *Aeromonas spp.* | *A. caviae* | *A. hydrophila/puntacta* | *A. caviae* |
| 22 | *V. fluviallis* | *A. caviae/ hydrophila* | *A. hydrophila/puntacta* | *A. sanarelli* |
| 23 | *Aeromonas spp.* | *A. caviae/ hydrophila* | *A. hydrophila/puntacta* | *A. caviae* |
| 24 | *Aeromonas spp.* | *A. caviae* | *A. hydrophila/puntacta* | *A. caviae* |
| 25 | *Aeromonas spp.* | *A. caviae* | *A. hydrophila/puntacta* | *A. caviae* |
| 26 | *Aeromonas spp.* | *A. caviae/ hydrophila* | *A. hydrophila/puntacta/sobria* | *A. sanarelli* |
| 27 | *Aeromonas spp.* | *A. caviae* | *A. hydrophila/puntacta* | *A. caviae* |
| 28 | *Aeromonas spp.* | *A. hydrophila* | *A. hydrophila/puntacta* | *A. hydrophila* |

**FIG 2.**
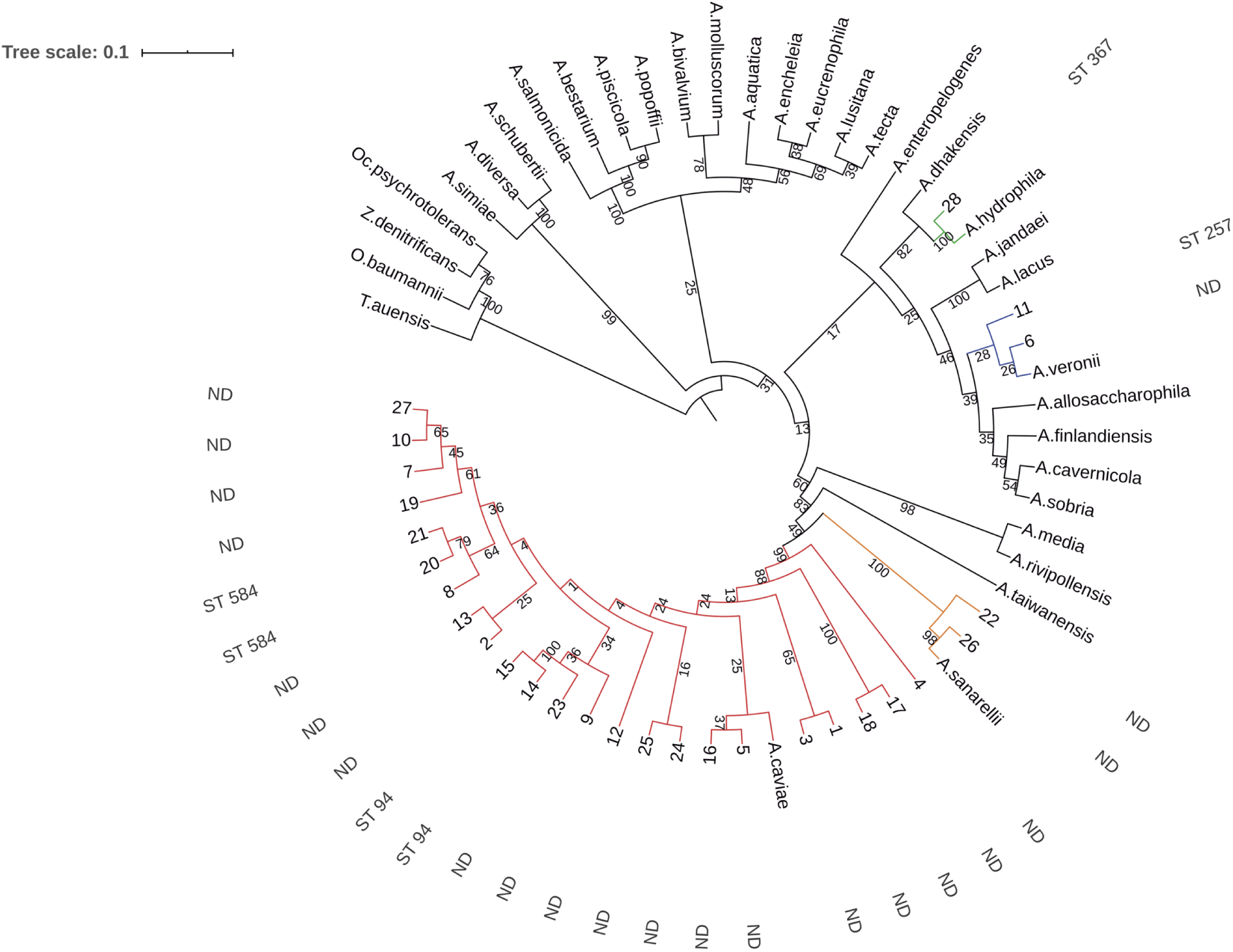
Phylogenetic maximum parsimony tree obtained from the concatenated sequences of seven housekeeping genes (*gyr*B, *gro*L, *glt*A, *met*G, *pps*A, *rec*A, and *rpo*B) showing the relationships between *Aeromonas* species, other genera of family *Aeromonadaceae*, such as *Tolumonas* (*T. auensis*), *Zobellella* (*Z. denitrificans*), *Oceanimonas* (*O. baumanii*), *Oceanisphaera* (*O. psychrotolerans*), and isolates indicated by Arabic numbers (1 to 28). Numbers at nodes indicate bootstrap values of percentage calculated using 2,000 replicates. The results obtained for MLST are shown as external values in the tree, ND stands for “ST not defined”.

## DISCUSSION

This study reports new insights regarding the repertoire of antimicrobial resistance genes of *Aeromonas* spp. isolated from diverse aquatic microbiomes, such as healthcare and domestic effluents, in addition to river water (Figure 1). Although the expression of β lactamases by *Aeromonas* spp. had been reported previously, the present study adds to the existing body of knowledge by carefully analyzing the genetic determinants of resistance in 28 unique cefotaxime-resistant *Aeromonas* isolates from a particular area of Brazil.

In our study, MDR *Aeromonas* spp. harboring GES-type carbapenemase were found in the hospital effluent, which was expected due to the use of large amounts of antimicrobials to prevent and treat infections. As that effluent does not receive any disinfecting pretreatment, this may contribute to the dissemination of MDR bacteria that were also found in the sanitary effluent located near the Complexo Hospital de Clínicas (Universidade Federal do Paraná) (CHC-UFPR). Our results are consistent with previous studies, showing that MDR *Aeromonas* strains are spreading rapidly in different environmental compartments and may be involved in the maintenance and dissemination of carbapenemase genes (14, 18, 29–31).

Despite several studies reporting the emergence of MDR and/or β-lactamase-producing *Aeromonas* isolates (30, 31) and rare reports of *bla*_KPC-2_ (17, 29, 32, 33), rare β-lactamase-encoding genes identified in our study deserve further attention. In particular, *bla*_GES-1_ and *bla*_GES-7_ encode ESBLs, whereas *bla*_GES-5_ and *bla*_GES-16_ encode enzymes able to hydrolyze carbapenems (34). GES-1 has been occasionally found in environmental samples, e.g., in a *Citrobacter freundii* strain (35). In our study, we detected GES-1 in two *A. caviae* isolates recovered from the Pequeno Principe Hospital (HPP) effluent. GES-7 has already been reported in *A. veronii* from river Seine in France (36) and *Aeromonas* spp. from WWTP in Poland (30). In our study, this enzyme was found in an MDR *A. veronii* from HPP effluent. This isolate was resistant to cephalosporins, carbapenems, aminoglycosides, and quinolones (Table 1). The resistance to carbapenems can be explained by the expression of the *cphA* chromosomal gene that may be species-specific, as it has been reported mainly in *A. hydrophila, A. veronii, A. jandaei*, and *A. dhakensis* (37–39).

GES-type carbapenemase producers, such as GES-5, GES-16, GES-24, and GES-31, have been recently reported in *Aeromonas* spp. from Brazil and Japan (18, 40, 41). Despite their importance for public health and increased incidence in healthcare facilities (42), the knowledge about carbapenem-resistant bacteria and circulation of their genes in the environment is very limited. GES-16, which differs from GES-5 by a single amino acid substitution, was first described in two *Serratia marcescens* clinical isolates from Rio de Janeiro, Brazil (43). These two GES-variants were recently described in *Aeromonas* spp. recovered from sea water also in Brazil (18). We found *bla*_GES-16_ in one *A. veronii* strain recovered from the HPP effluent and *bla*_GES-5_ in one *A. caviae* recovered from the CHC-UFPR effluent (Table 1). These findings are alarming because *bla*_GES_ genes are essentially gene cassettes associated with integrons on plasmids that increase the risk of rapid horizontal gene transfer and interspecies dissemination of virulence and resistance determinants (34, 44).

Growing levels of drug resistance, especially to β-lactam, have been reported in *Aeromonas* spp. not only in clinical isolates, but also in isolates from water ecosystems and aquatic organisms (13, 29, 31). In our study, CTX-M β-lactamase harboring strains were the prevalent group in WWTP that receives high amounts of microbial contaminants from hospitals effluents and other sources as well as in DRW. Such strains are globally endemic in clinical settings and represent most frequent sources of ESBLs in community-acquired infections and livestock (45–48). Few studies have reported the presence of *bla*_CTX-M_ genes in the environmental *Aeromonas* spp. (29–31, 49–52) and as far as we know, this is the first study reporting environmental *A. caviae* that expresses both *bla*_CTX-M-2_ and *bla*_CTX-M-9_.

ESBL-producing bacteria often present co-resistance to other antimicrobials, particularly ciprofloxacin (53–56). Among the PMQR genes screened for, only *aac(6′)-Ib-cr* and *qnr*S were detected, often in combination, in the species *A. caviae, A. veronii*, and *A. hydrophila. aac(6′)-ib-cr* and *qnr*S were detected, respectively, in six and three isolates from the HPP and sanitary effluents (Table 1). These results are in agreement with previous reports about the widespread presence of quinolone resistance genes in the environment and clinical settings, as well as their significant prevalence in hospital effluents compared to that in municipal wastewater (57–60). This could be explained by the use of quinolones in clinical practice and by the persistence in the environment of the active form of ciprofloxacin excreted with feces and urine (61, 62). Although these genes confer low-levels of resistance, their presence can favor and complement the selection of other resistance mechanisms (63).

The knowledge of the main characteristics of *Aeromonas* species and strains, such as ecological, environmental, and host distributions, is currently hampered by the lack of precise delineation of genetic clusters at the species, subspecies, and clone levels. There are few studies about ecological niches for different STs of *Aeromonas* spp. (3, 64). In our study, we accurately assigned ST 257 to *A. veronii* and ST 94 to *A. caviae* (Figure 2), whereas in PubMLST database, the latter corresponds to *A. hydrophila*. Taking into account the period when that reference was deposited (2011), the methods for identifying *Aeromonas* species were still under development and species were assigned as *A. hydrophila*/*A. caviae*. The major problem of the MLST scheme that was created in 2010 (25) is that the comparisons might be limited by the number of strains and origins available in the database at that time. The last update of the database (06 May 2020) had 755 strains and 3,038 sequences that corresponded to 689 MLST profiles (https://pubmlst.org/Aeromonas/submission.shtml, accessed on 21 May 2020) (65).

There was an agreement between the results of MLST and MLPA approaches. Phylogenetic tree analysis classified 28 *Aeromonas* isolates into four different species, with *A. caviae* being the most prevalent. Other studies also demonstrated that *A. caviae* is in fact the predominant species in polluted environments (66–69). This species along with *A. veronii* and *A. hydrophila*, also detected in our study, are responsible for causing bacteremia, gastroenteritis, or even septicemia in both immunocompromised and immunocompetent individuals (58). These isolates were previously identified by MALDI-TOF with 100% agreement at the genus level, and with 92.9% agreement at the species level. These results are relatively similar to those reported by other authors (70, 71), suggesting that MALDI-TOF is a useful tool, because the identification error was <10%. Overall, housekeeping gene sequencing with phylogenetic analysis was found to be the most accurate in identifying *Aeromonas* at the species level.

The finding of MDR strains, almost all in the hospital effluent, and other determinants of resistance in all collected sites surviving conventional treatments emphasizes the notion that *Aeromonas* spp. could serve as a vehicle and an important reservoir of ARG and reinforces the need for effective actions to contain the spread of ARG and antibiotic resistant bacteria from the environment to human and animal niches. A major problem in controlling these spread in the environment is the lack of national or international legal regulations (72).

Our study serves the objective of prevention and control of antimicrobial resistance (AMR), which is treated in the global and national context, respecting the One Health approach, which requires the joint work of human, animal, and environmental health. National Action Plan for the Prevention and Control of Antimicrobial Resistance (PAN-BR) (73) was developed in convergence with the objectives defined by the tripartite alliance between the World Health Organization (WHO), the United Nations Food and Agriculture Organization (FAO) and the World Organization for Animal Health (OIE) and presented in the Global Action Plan on Antimicrobial Resistance (74). Our study strengthens the knowledge and scientific basis regarding the spread of antimicrobial resistance through *Aeromonas* spp. recovered from aquatic environments.

The identification of the microorganisms that may be responsible for inter- and intra-species transmission of genes conferring antimicrobial resistance in relation to their ecological habitats is an important line of microbiological research. Phylogenetic analyses based on MLST/MLPA proved to be a valid and practical method for species identification. However, the insufficient adequacy of the currently available public resource for sequence typing of *Aeromonas* strains is an issue that may require substantial improvement of the PubMLST database. The taxonomy of this genus is complex due to high identity among nucleotide sequences of different species. The database should include newly described species as well as undergo reclassification, taking into account amended or extended descriptions of existing taxa.

## MATERIALS AND METHODS

### Sample collection and bacterial isolates

A single wastewater sample was collected from different aquatic environments located in Curitiba, Brazil. Figure 1 shows schematic representation of different sample collection sites.

Samples were collected into sterile 1 L sampling bottles and processed by the membrane filtration method as previously described (75). Oxidase- and glucose-positive single colonies were selected from MacConkey agar plates supplemented with cefotaxime (2 mg/L).

Bacterial species were identified by VITEK^®^ 2 Compact System (Biomérieux, Marcy-l’Etoile, France) and mass spectrometry using Microflex LT Biotyper 3.0 (Bruker Daltonics, Bremen, Germany) and VITEK^®^ MS (Biomérieux, Marcy-l’Etoile, France) instruments. To select distinct genetic profiles, ERIC-PCR was performed using primers and amplification conditions described in Supplementary Table 1.

### Antimicrobial resistance characterization

Antimicrobial susceptibility testing was performed using agar dilution according to the Clinical and Laboratory Standards Institute method (76). Double disc synergy was assessed to detect ESBL (77), class A and B carbapenemases (78).

To identify β-lactamase genes (*bla*_TEM_, *bla*_SHV_, *bla*_CTX-M,_ *bla*_GES_, *bla*_PER_, *bla*_KPC_, *bla*_NDM_, *bla*_VIM_, *bla*_SIM_, *bla*_GIM,_ *bla*_BES_, *bla*_VEB,_ *bla*_SPM,_ *bla*_IMP,_ and *bla*_OXA_), PMQR targets (*qnr*A, *qnr*B, *qnr*C, *qnr*D, *qnr*S, *qnr*VC, *qep*A, *oqx*AB, and *aac(6′)-Ib-cr*), and 16S RMTases (*npm*A, *arm*A, and *rmt*A–H), PCR was performed using primers and amplification conditions indicated in Supplementary Table 1.

PCR products were sequenced using a 3730XL DNA Analyzer (Applied Biosystems, Carlsbad, CA, USA). Nucleotide and protein sequences were analyzed using Lasergene Software Package (DNASTAR, Madison, WI, USA). Resistance gene references were selected in GenBank database.

### MLST and phylogenetic analysis

MLST was performed by PCR and sequencing of six *Aeromonas* spp. housekeeping genes (*gyr*B, *gro*L, *glt*A, *met*G, *pps*A, *rec*A) according to Martino et al. (25) and classified by using web-based MLST sequence database (http://pubmlst.org/aeromonas, accessed on March 2020).

For MLPA, multiple alignments containing the concatenated sequences were aligned in the following gene order *gyr*B-*gro*L‐*glt*A‐*met*G‐*pps*A‐*rec*A, starting and ending at exactly the same positions. The *rpo*B gene was added to the sequence of these genes to build a more reliable phylogenetic tree (79). The phylogenetic tree included 29 representative *Aeromonas* spp. and other genera of the family Aeromonadaceae: *Tolumonas* (*T. auensis*), *Zobellella* (*Z. denitrificans*), *Oceanimonas* (*O. baumanii*), and *Oceanisphaera* (*O. psychrotolerans*). Twelve species were not included in the MLPA analysis because some of their housekeeping genes were not available in GenBank databases (Figure 2). Seaview 4 software (80) was used to apply the phylogenetic method of maximum parsimony (81) with Bootstrap values calculated using 2,000 replicates as default. The phylogenetic tree was visualized by Interactive Tree Of Life (v.4) (82) available at https://itol.embl.de/.

## ACKNOWLEDGMENTS

We thank Central Laboratory of Paraná (LACEN), Paraná, Brazil for performing the MALDI-TOF assay and the staff of the Life Sciences Core Facility (GoGENETIC) from Federal University of Paraná (UFPR) for DNA sequencing. We also thank the visual designer Ricardo Hurmus for helping with the graphical illustration of sample collection sites.

